# Membrane lipid requirements of the lysine transporter Lyp1 from *Saccharomyces cerevisiae*

**DOI:** 10.1101/2020.03.27.011031

**Authors:** Joury S van‘t Klooster, Tan-Yun Cheng, Hendrik R Sikkema, Aike Jeucken, D. Branch Moody, Bert Poolman

## Abstract

Membrane lipids act as solvents and functional cofactors for integral membrane proteins. The yeast plasma membrane is unusual in that it may have a high lipid order, which coincides with low passive permeability for small molecules and a slow lateral diffusion of proteins. Yet, membrane proteins whose functions require altered conformation must have flexibility within membranes. We have determined the molecular composition of yeast plasma membrane lipids located within a defined diameter of model proteins, including the APC-superfamily lysine transporter Lyp1. We now use the composition of lipids that naturally surround Lyp1 to guide testing of lipids that support the normal functioning of the transporter, when reconstituted in vesicles of defined lipid composition. We find that phosphatidylserine and ergosterol are essential for Lyp1 function, and the transport activity displays a sigmoidal relationship with the concentration of these lipids. Non-bilayer lipids stimulate transport activity, but different types are interchangeable. Remarkably, Lyp1 requires a relatively high fraction of lipids with one or more unsaturated acyl chains. The transport data and predictions of the periprotein lipidome of Lyp1, support a new model in which a narrow band of lipids immediately surrounding the transmembrane stalk of a model protein allows conformational changes in the protein.

## Introduction

In the fluid mosaic bilayer model(1) lipids are seen as solvent for integral membrane proteins like aqueous media are for soluble proteins. The length of the lipids should approximately match the hydrophobic region of the protein, and the polar headgroups of the lipids can interact with the hydrophilic regions of the membrane protein(2). These parameters are referred to as hydrophobic-hydrophilic matching of proteins and lipids(3), which has been shown to affect the activity and or stability of membrane proteins(4–6). The degree of saturation of acyl chains influence the packing of lipids and thus the fluidity of the bilayer. Additionally, sterols can increase the order of the acyl chains(7), which is accompanied by a decrease of the lateral diffusion of components in the membrane(8). Sphingolipids typically have saturated long-chain fatty acids and preferentially interact with sterols, giving rise to so-called liquid-ordered membrane domains that are phase-separated from more fluid domains(9). The hydrophilic heads of phosphatidylglycerol (PG) and - serine (PS) lipids form hydrogen bonds with polar amino acids, and salt-bridges can be formed between basic residues on the protein and the lipid phosphate(10).

In addition to specific bonding between lipids and proteins, the geometry of lipids can influence the structure and hence the activity of proteins through their impact on the physical properties of the membrane(11). Lipids with a cylindrical shape, like phosphatidylcholine (PC), PG and PS, have a head-group cross-sectional area comparable to that of the acyl chains and spontaneously form bilayers. Lipids with a cone shape, like phosphatidylethanolamine (PE) and phosphatidic acid (PA), have a head-group cross-sectional area that is smaller than that of the acyl chain area. These so-called non-bilayer lipids are capable of forming a hexagonal phase and induce substantial negative membrane curvature(12). The different geometry of bilayer and non-bilayer types of lipids translate into differences in the lateral pressure profile of the membrane that in turn can modulate protein conformations(13). The lateral pressure profile is also affected by the acyl chain length and degree of unsaturation, and charge of the headgroups(14).

The lipids directly surrounding the membrane protein are referred to as annular lipids. High-resolution X-ray(15–17) and cryo-TEM structures(18) and electron-spray-ionization mass spectrometry (ESI-MS)(19–22) of membrane proteins have revealed non-annular ‘structural lipids’ that specifically interact in between transmembrane α-helices or in hydrophobic pockets of transmembrane domains. Thus, lipid-protein interactions are likely crucial in the activity of the majority of membrane proteins but actual functional data to support this hypothesis is scarce. The picture that emerges from studies on four different bacterial transporters (23–26) is that anionic lipids can fulfill specific roles in the activity or and regulation of transport activity, and that non-bilayer lipids stimulate transport. The role of lipids in the conformational dynamics of bacterial transporters has also been reported(27–29). However, we are not aware of systematic studies on the function of lipids in eukaryotic plasma membrane transporters, but the importance of the topic has been recognized(30–33).

In our recent paper (eLIFE, revision under review), we used SMALPs to isolate lipids located within a defined diameter of the transmembrane stalk of proteins in the plasma membrane of yeast. Such ‘periprotein lipids’ form in a stoichiometric excess of ~60:1 with the protein. They encompass more than the annular lipids and represent additional shell(s) of lipids surrounding the protein. We find that the Lyp1 periprotein lipidome is enriched in phosphatidylserine and depleted in ergosterol relative to the overall composition of the plasma membrane. Using these data and the availability of synthetic lipids in defined vesicles as starting point, we systematically analyzed the lipid dependence of transport of the eukaryotic amino acid proton-symporter Lyp1 from *Saccharomyces cerevisiae*.

## Results and Discussion

### Phospholipid dependence of Lyp1 function

We previously reconstituted Lyp1 in lipid vesicles composed of yeast total lipid extract(34), and we now incorporated the protein into vesicles with a defined lipid composition. The yeast sphingolipids and certain phospholipids with precisely matching headgroup-acyl chain combinations identified in our periprotein lipidome analyses are not available, but we could design lipid mixtures that otherwise mimic periprotein lipids in yeast cells. To drive the import of lysine by Lyp1, we impose a membrane potential (∆Ψ) and pH gradient (∆pH) by diluting the vesicles containing K-acetate into Na-phosphate plus valinomycin. The driving force is determined by the potassium and acetate gradient (Fig. 1A). Typically, we use a ∆Ψ and -Z∆pH of −81 mV each (Z=58 mV), producing a proton motive force of −162 mV. We compared the lipid composition of the Lyp1 vesicles with the starting mixture for the reconstitution, using mass spectrometry, and did not find significant differences (Fig. S1). This rules out enrichment or depletion of certain lipids during membrane reconstitution, including the possibility that co-purified protein lipids contribute significantly to the lipid pool. Furthermore, proton permeability measurements indicate that the transport rates are not skewed by large differences in proton leakage (Fig. S2). Hence, the proton motive force is similar in vesicles of different lipid composition. Figure 1B shows the import of lysine over time, from which the slope estimates the initial rate of transport.

**Figure 1:**
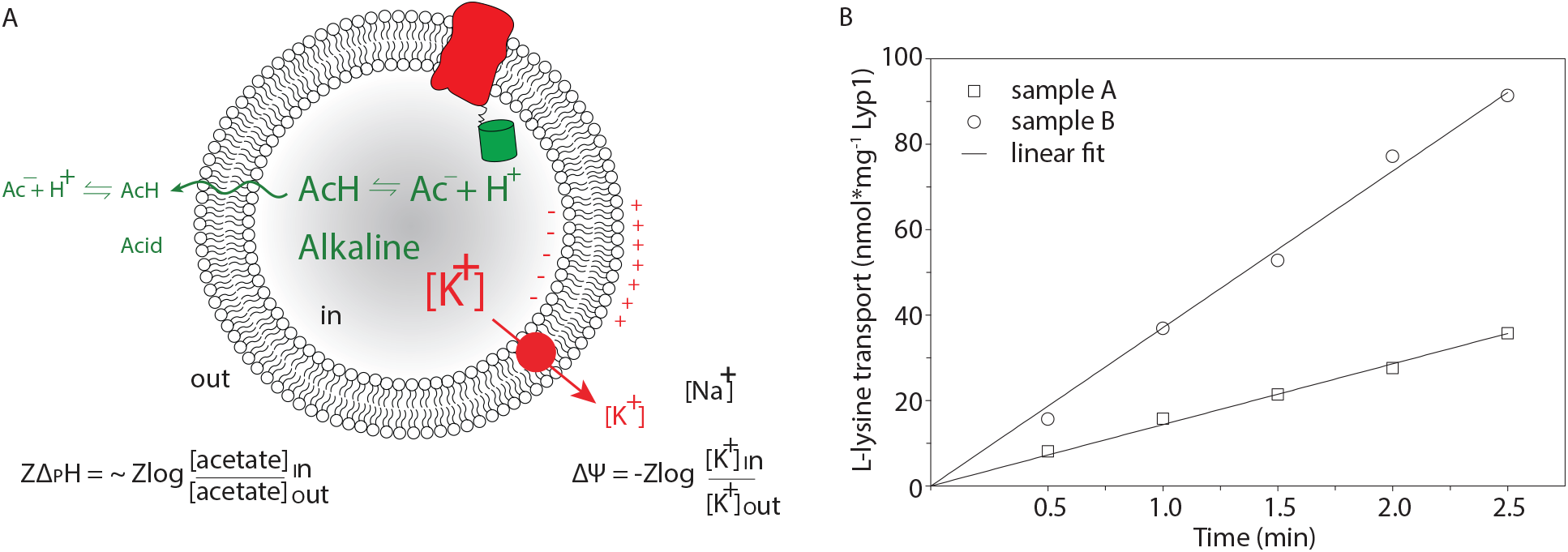
Generation of proton motive force and lysine transport progress curves. (A) Schematic showing the generation of a membrane potential (ΔΨ, red arrow) by a valinomycin-mediated potassium diffusion potential and pH gradient (ΔpH, green) via an acetate diffusion potential. Together the ΔΨ and ΔpH form the proton motive force (PMF=ΔΨ-ZΔpH, where Z equals 2.3RT/F and R and F are the gas and Faraday constant, respectively, and T is the absolute temperature. (B) Transport of lysine by Lyp1-GFP-containing proteoliposomes and data fitting. Lyp1 activity is obtained from the slopes of such lines and the rates of transport are converted into turnover numbers

In initial experiments with ergosterol present we observed that C34:1 (C18:1 plus C16:0 acyl chains) palmitoyl-oleoyl-*sn*-phosphatidylX (POPX, where X=choline, ethanolamine, glycerol or serine; Figure 2A-B), support a much higher transport activity than the most commonly used C36:2 forms or when C32:0 dipalmitoyl-*sn*-phosphatidylX (DPPX; C32:0 = 2 C16:0 chains) lipids were used (Fig. 2C). Hence, we started with a mixture of POPS, POPG, POPE and POPC (17.5mol% each) plus 30mol% ergosterol (Fig. 2D, #1) and used that to benchmark the activity of Lyp1 in vesicles against the effects of phospholipids and sterols. Figure 2D shows that increasing either POPS or POPE increases transport activity (#2 and #3), while reducing the fraction of these lipids decreases the activity (#5 and #6) relative to that in the benchmark mixture (#1). Increasing the fraction of POPG or lowering of POPC (#4) had no negative effect on transport. This suggests that the non-bilayer lipid POPE and the anionic lipid POPS are a minimal requirement for transport (#7).

**Figure 2:**
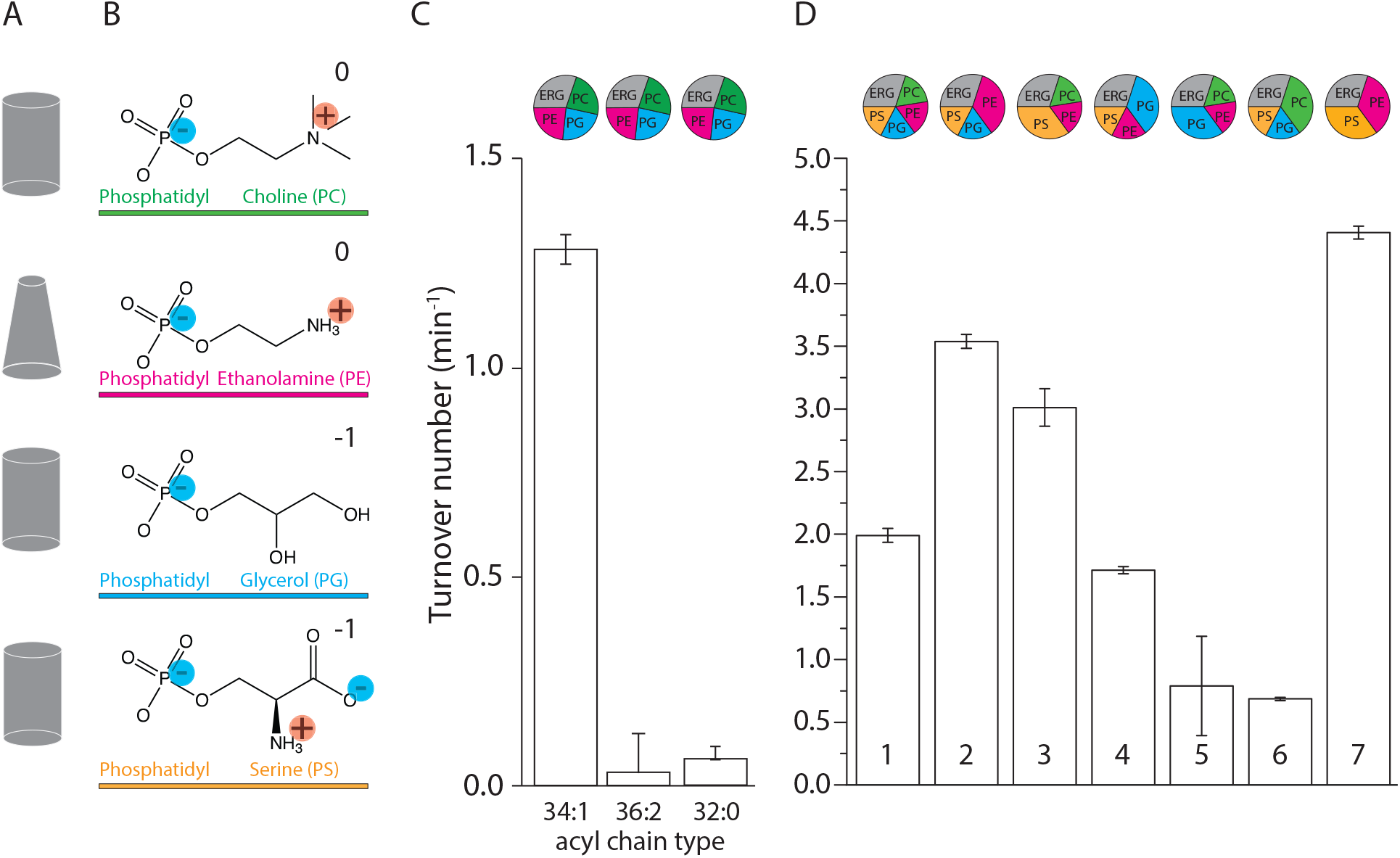
Lyp1 activity as a function of lipid composition. (A) shows the geometric representation of lipids for the headgroups shown in B. (B) shows the headgroups of phospholipids (and color-coding) with the net charge of the lipids at pH 7. (C) shows the turnover number of Lyp1 in lipid mixtures with different acyl chains. (D) shows the turnover number of Lyp1 in different lipid mixtures with C34:1 (PO) acyl chains. The lipid composition (mol%) of each sample is visualized by pie-graphs, using the color coding of (B). The quantity of ergosterol was kept at 30mol% in all mixtures. Data based on 3 replicate experiments; the bars show the standard error of the fit.

### The role of anionic lipids

PG and PS are both anionic lipids, therefore it is not surprising that PG can partly substitute PS (Fig. 2D, #5). Yet, PG is the main anionic lipid found in bacteria, while PS is the predominant anionic lipid in eukaryotes. It is thus likely that Lyp1 has evolved to function optimally in membranes containing PS. Next, we simplified the lipid mixture by preparing vesicles composed of POPS, POPE plus Ergosterol (Fig. 2D, #7) and step-wise reduced the quantity of PS. We found a sigmoidal relationship between Lyp1 activity and POPS concentration (Fig. 3A), which is indicative of cooperativity and suggests that more than one molecule of POPS is needed for activation of Lyp1.

**Figure 3:**
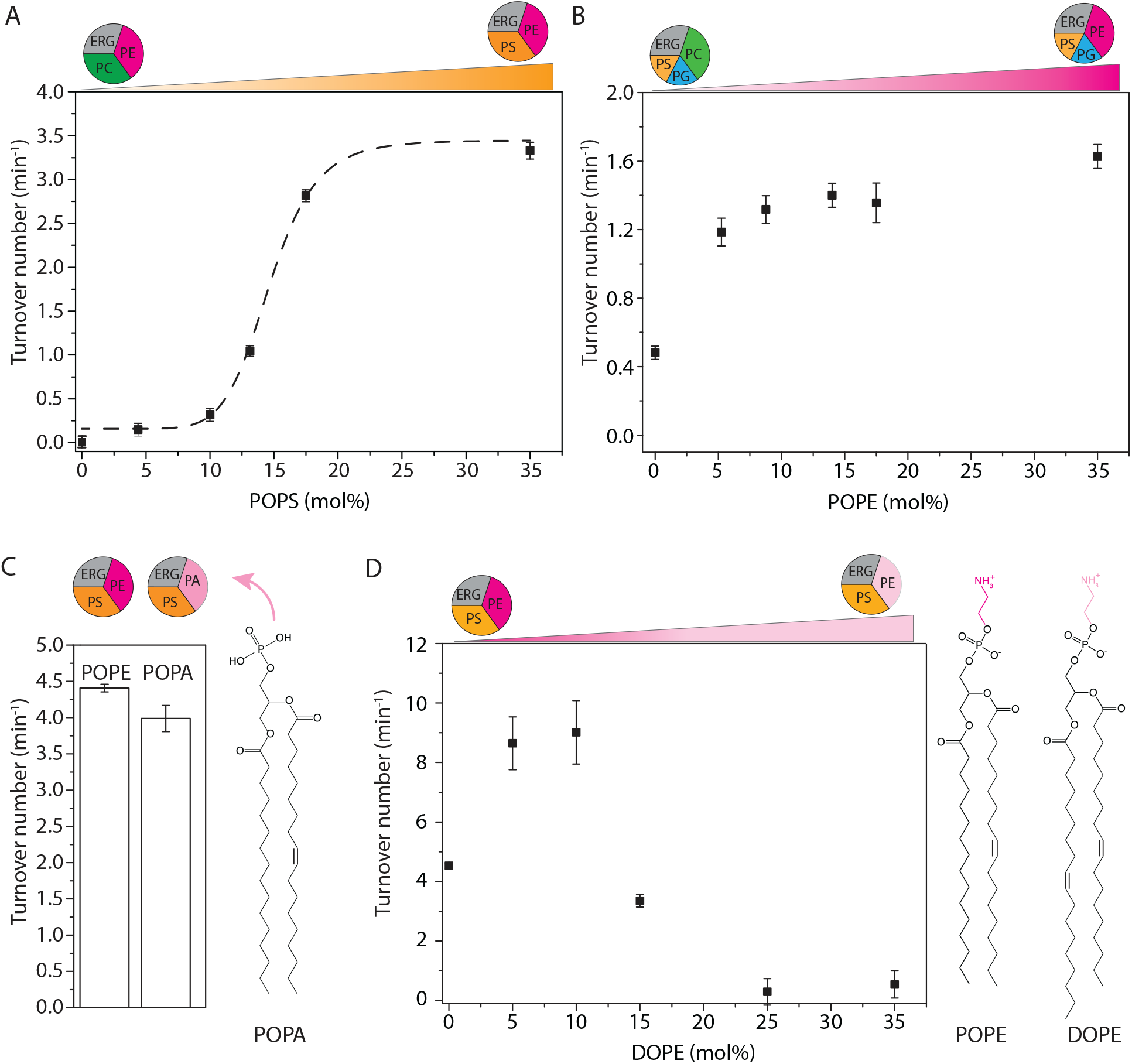
Effect of anionic and non-bilayer lipids on Lyp1 activity in proteoliposomes. (A) Turnover number of Lyp1 as a function of POPS (1-palmitoyl-2-oleoyl-*sn*-glycero-3-phosphoserine). (B) Turnover number of Lyp1 as a function of POPE (1-palmitoyl-2-oleoyl-*sn*-glycero-3-phosphoethanolamine) (mol%). (C) Turnover number of Lyp1 in vesicles with POPE versus POPA (1-palmitoyl-2-oleoyl-sn-glycero-3-phosphatidic-acid). (D) Turnover number of Lyp1 as a function of DOPE, which was increased at the expense of POPE. The triangles at the top of each graph depict the gradual replacement of one lipid for another. Number of replicate experiments (n) = 3; the variation between replicates is ± 20% and therefor the error bars are the standard error of the fit of n = 1. Sigmoidal curves were fitted using the equation: 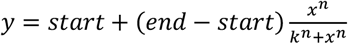

### The role of non-bilayer lipids

Many phospholipids have a cylindrical shape that allow bilayer formation with a single lipid species. The relatively small headgroup of PE, as compared to the acyl chains, results in a cone-shaped geometry that does not allow bilayer formation from pure PE. PE and other non-bilayer lipids affect the lateral pressure profile and membrane curvature more drastically than bilayer forming lipids do(35). We determined the PE dependence of Lyp1 by increasing the fraction of POPE at the expense of POPC (Fig. 3B). Lyp1 transport activity increases 3-fold with increasing POPE and saturates at ~10mol%. To determine whether the ethanolamine headgroup or the geometric shape of the lipid is important, we substituted POPE for POPA (Fig. 3C), a non-bilayer forming conical phospholipid devoid of the headgroup moiety in PE. POPA can fully substitute POPE in transporter activity, which suggests that PE is not essential but lipids with non-bilayer properties are important for Lyp1 function. Next, we titrated in DOPE at the expense of POPE to gradually increase the degree of acyl chain unsaturation (Fig. 3D). We observe a 2-fold increase in Lyp1 activity with DOPE at 5 to 10mol% and POPE at 30 to 25mol%, but a further increase in dioleoyl at the expense of palmitoyloleoyl chains decreases the activity to zero. These experiments indicate that specific features of the acyl chains, which affect membrane fluidity, are as important as the geometric shape of non-bilayer lipids.

### Ergosterol is essential for Lyp1 activity

Ergosterol is the major sterol of lower eukaryotes and present in the yeast plasma membrane at concentrations of ≈ 30mol%(36). We increased the fraction of ergosterol at the expense of POPC and observed an increase in activity up to 10mol%. Without ergosterol, Lyp1 is not active and the activity drops above 25mol% (Fig. 4A). As with POPS we find a sigmoidal dependence but the apparent cooperativity is much lower. The predominant sterol of mammalian cells, cholesterol(37), supports less than 15% of Lyp1 activity as compared to ergosterol. Ergosterol differs from cholesterol in that it has two additional double bonds at positions C7-8 and C22-23 and one extra methyl group at C24 (Fig. 4C yellow ovals). To find out which of these is important for transport activity, we tested brassicasterol and dehydrocholesterol and an equal mixture of both (Fig. 4C). However, the two sterols and the mixture thereof, which together represent the functional groups of ergosterol, cannot substitute for ergosterol itself (Fig 4B).

**Figure 4:**
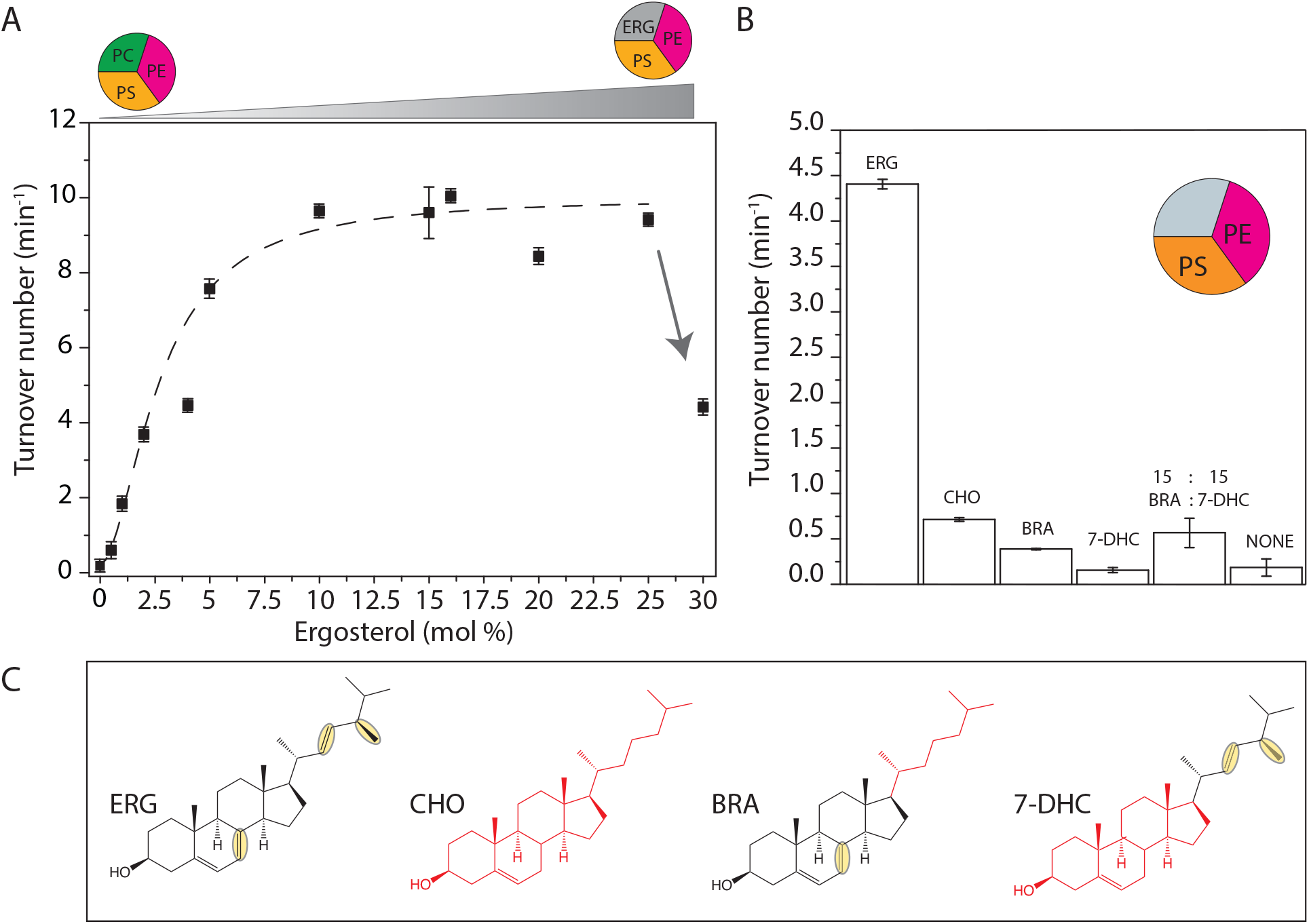
Effect of sterols on Lyp1 activity. (A) Turnover number of Lyp1 as a function of the quantity of ergosterol in the vesicles. Arrow indicates the drop off activity from 25 to 30mol% ergosterol. (B) Turnover number of Lyp1 as a function of the type of sterol. (C) Structures of sterols; ERG = Ergosterol; CHO = Cholesterol; BRA = Brassicasterol; 7-DHC = 7-Dehydrocholesterol. ERG and CHO parts are black and red respectively. Structural dissimilarities between ERG and CHO are highlighted by yellow ovals. Number of biological experiments (n) = 3; the variation between replicates is ± 20% and therefor the error bars are the standard error of the fit of n = 1.

### In conclusion

Our periprotein lipidomics study (eLIFE, revision under review) combined with published literature led to a new testable model of how proteins may function in a membrane of high lipid order(38–41). These biophysical properties coincide with the high tolerance of yeast for a low pH and high concentration of solvents, which allows the organism to survive in adverse environments(42,43).

We find that anionic lipids with saturated and unsaturated acyl chains, preferably POPS, and ergosterol are essential for Lyp1 functioning, and both lipid species stimulate transport cooperatively. This suggests that “allosteric” sites on the protein need to be occupied by specific lipids to enable transport. About 20mol% PS and 5mol% ergosterol are sufficient for optimal activity of Lyp1, which coincides with our periprotein lipidomic analysis (17-20mol% PS and 4-5mol% ergosterol). This is distinct from that of the total plasma membrane where PS and ergosterol amount to 6 and 30mol%, respectively, and this would reduce the apparent activity of Lyp1 severely. Hence, the periprotein lipidome provides an environment of specific lipids and sufficient local membrane fluidity for optimal functioning of Lyp1.

## Methods

### Cultivation of P. pastoris, protein expression and plasma membrane isolation

The expression of Lyp1 in *Pichia pastoris* (SMD1163-Lyp1-TEV-GFP-his_10_) and its membrane isolation was described previously (34). *P. pastoris* cultures were harvested by centrifugation at 7,500 × g, for 15 min at 4°C (Beckman centrifuge J-20-XP, Rotor JLA 9.1000, US). Further steps were performed at 4°C or on ice. The resulting cell pellet was resuspended in 150 mL of cell resuspension buffer (CRB; 20mM Tris-HCl pH6.7, 1mM EGTA, 0.6M sorbitol, 10μM pepstatin A (Apollo Scientific, UK), 10μM E-64 (Apollo Scientific, UK) plus protease inhibitors (cOmplete Mini EDTA-free™, ROCHE, 1 tablet/75 mL). Centrifugation was repeated and the resulting cell pellet was resuspended to OD_600_ = 200 in CRB with 1 tablet cOmplete Mini EDTA-free™/10 mL). The resulting cell suspension was broken using a cell disrupter (Constant cell disruption systems, US) by three sequential passes operating at 39Kpsi, after which 2 mM fresh PMSF was added. Unbroken cells were pelleted by centrifugation at 18,000 RCF for 30 min at 4°C (Beckman centrifuge J-20-XP, Rotor JA16.250, DE). The supernatant was transferred to 45 TI rotor (Beckman, DE) tubes and membranes were pelleted by ultracentrifugation (Optima XE-90, Beckman, DE) at 186,000 RCF, 90 min at 4°C. The resulting membrane pellet was resuspended to homogeneity at 400 mg/mL using a potter Elvehjem tissue grinder in 20 mM Tris-HCl pH7.5, 0.3M Sucrose, 10μM Pepstatin A, 10μM E-64 (Apollo Scientific, UK), 2 mM PMSF and protease inhibitor cocktail (cOmplete Mini EDTA-free™, 1 tablet/10 mL). Aliquots of 2 mL were snap frozen using liquid nitrogen and stored at −80°C until further use.

### Liposome formation

Phospholipids were obtained from Avanti polar lipids Inc (Alabaster, AL, USA). Brassicasterol (CarboSynth, UK), 7-dehydrocholesterol (Sigma Aldrich, DE), cholesterol and ergosterol (Sigma Aldrich, DE) were obtained from the indicated vendor. Lipids were dissolved in chloroform and mixed at desired ratios (mol%) to a total weight of 10 mg in a 5 mL glass round bottom flask. Chloroform was removed by evaporation at 40°C and an applied pressure of 370 mBar, using a rotary vaporizer (rotavapor r-3-BUCHI). The resulting lipid film was resuspended in 1 mL of diethylether and the previous step was repeated at atmospheric pressure until a dry film was visible. To remove any residual solvent a pressure of 10 mBar was applied for 20 minutes. The obtained lipid film was hydrated in 1mL of 50 mM NH_4_-acetate pH7.5, 50 mM NaCl by shaking for 5 min and then transferred to a plastic tube compatible with sonication. The lipid suspension was homogenized by tip sonication with a Sonics Vibra Cell sonicator (Sonics & Materials Inc.) at amplitude of 70% for 2 minutes with 5 sec pulses and 5 sec pauses between each pulse. The sample was kept at 4°C in ethanol:water:ice (25:25:50 v/v/v). The lipid suspension (1 mL aliquot at 10 mg of lipid/mL) was transferred to a 1.5 mL Eppendorf tube, flash frozen in liquid nitrogen and thawed at 40 °C. This process was repeated 4 times and stored in liquid nitrogen until further use.

### Proteoliposome preparation

Protein purification and membrane reconstitution was performed as described (34). Briefly, *n*-Dodecylmaltoside (DDM) was used to solubilize Lyp1-GFP. The protein was purified by Immobilized Metal Affinity Chromatography (IMAC, using Nickel-Sepharose) and Fluorescence Size-Exclusion Chromatography (using a Superdex 200 increase 300/10 GL column attached to an Åkta 900 chromatography system (Amersham Bioscience, SE) with in-line Fluorescence detector (1260 Infinity, Agilent technologies, US). In parallel, liposomes were thawed at room temperature and subsequently homogenized by 11 extrusions through a 400nm polycarbonate filter (Avestin Europe GMBH, Ger). Next, liposomes were destabilized using Triton X-100 and titrated to a point beyond the saturation point (Rsat) as described(45); the final turbidity at 540 nm was at approximately 60% of Rsat. Purified Lyp1 was mixed with triton x-100-destabilized liposomes of the appropriate lipid composition at a protein-to-lipid ratio of 1:400 and incubated for 15 min under slow agitation at 4°C. Bio-beads SM-200 (Biorad, Hercules, Ca, USA), 100 mg/0.4% Triton X-100 (final concentration of Triton x-100 after solubilization of the liposomes) were sequentially added at 15, 30 and 60 min. The final mixture was incubated overnight, after which a final batch of bio-beads was added and incubation continued for another 2 hours. Protein containing liposomes (proteo-liposomes) were separated from the Bio-beads by filtration on a column followed by ultracentrifugation 444,000 × g at 4°C for 35 min. Proteo-liposomes were suspended in 10 mM potassium-phosphate plus 100 mM potassium-acetate pH 6.0, snap frozen and stored in liquid nitrogen.

### In vitro transport assays

Transport assays were performed and the formation of a proton gradient (∆pH) and membrane potential (∆Ψ) was established as described (34). Both ∆pH and ∆Ψ were formed by diluting the proteo-liposomes (suspended in 10 mM potassium-phosphate plus 100 mM potassium-acetate pH 6.0) 25-fold into 110 mM sodium-phosphate pH6.0 plus 0.5 μM valinomycin. This results in maximal values of Z∆pH and ∆Ψ of −83 mV as calculated according to the Nernst equation, yielding a proton motive force of −166 mV.

### Data analysis and transport rates

The data were analyzed in Origin (OriginLab, MA). The initial rates of transport were determined from the slope of progress curves (Figure 1B). The relative transport activity was determined by normalizing each vesicle preparation to the sample with a lipid composition POPC:POPE:POPS:POPG:Ergosterol in a ratio of 17.5:17.5:17.5:17.5:30 (mol%).

### Lipid extraction and mass spectrometry

Lipid extraction from the proteo-liposomes was performed based on the Bligh and Dyer method (46). The lower organic phase was separated from the upper aqueous phase and dried under a nitrogen stream. The extracted lipid residue was re-dissolved in the starting mobile phase A and 10ul of the lipid extraction was injected on a hydrophilic interaction liquid chromatography (HILIC) column (2.6 μm HILIC 100 Å, 50 × 4.6 mm, Phenomenex, Torrance, CA). The mobile phases were (A) acetonitrile/acetone (9:1, v/v), 0.1% formic acid and (B) acetonitrile/H2O (7:3, v/v), 10mM ammonium formate, 0.1% formic acid. In a 6.5-minute run, the solvent gradient changes as follows: 0-1 min, from 100% A to 50% A and B; 1-3 min, stay 50% B; 3-3.1 min, 100% B; 3.1-4 min, stay 100% B; 4-6.5 min from 100% B to 100% A. Flowrate was 1.0mL/min The column outlet of the LC system (Accela/Surveyor; Thermo) was connected to a heated electrospray ionization (HESI) source of mass spectrometer (LTQ Orbitrap XL; Thermo) operated in negative mode. Capillary temperature was set to 350°C, and the ionization voltage to 4.0 kV. High resolution spectra were collected with the Orbitrap from m/z 350-1750 at resolution 60,000 (1.15 scans/second). After conversion to mzXML data were analyzed using XCMS version 3.6.1 running under R version 3.6.1.

## Acknowledgements

This work was carried out within the BE-Basic R&D Program, which was granted a FES subsidy from the Dutch Ministry of Economic affairs, agriculture and innovation (EL&I), and was supported by an ERC Advanced Grant (ABCVolume; #670578). The research was also funded by NWO TOP-PUNT (project number 13.006) grants and the National Instiutes of Health (R01 AR048632).

## Author contributions

J.S.v.t.K., D.B.M. and B.P designed the research plan; J.S.v.t.K performed the research except for the liquid-chromatography coupled to mass spectrometry analysis; J.S.v.t.K., T-Y.C., D.B.M. and B.P. analyzed the data; A. Jeucken performed mass spectrometry analysis corresponding to figure S1; J.S.v.t.K., T-Y.C., D.B.M. and B.P. wrote the paper.

## SUPPLEMENTARY INFORMATION

### Supplementary figures

**Figure S1:**
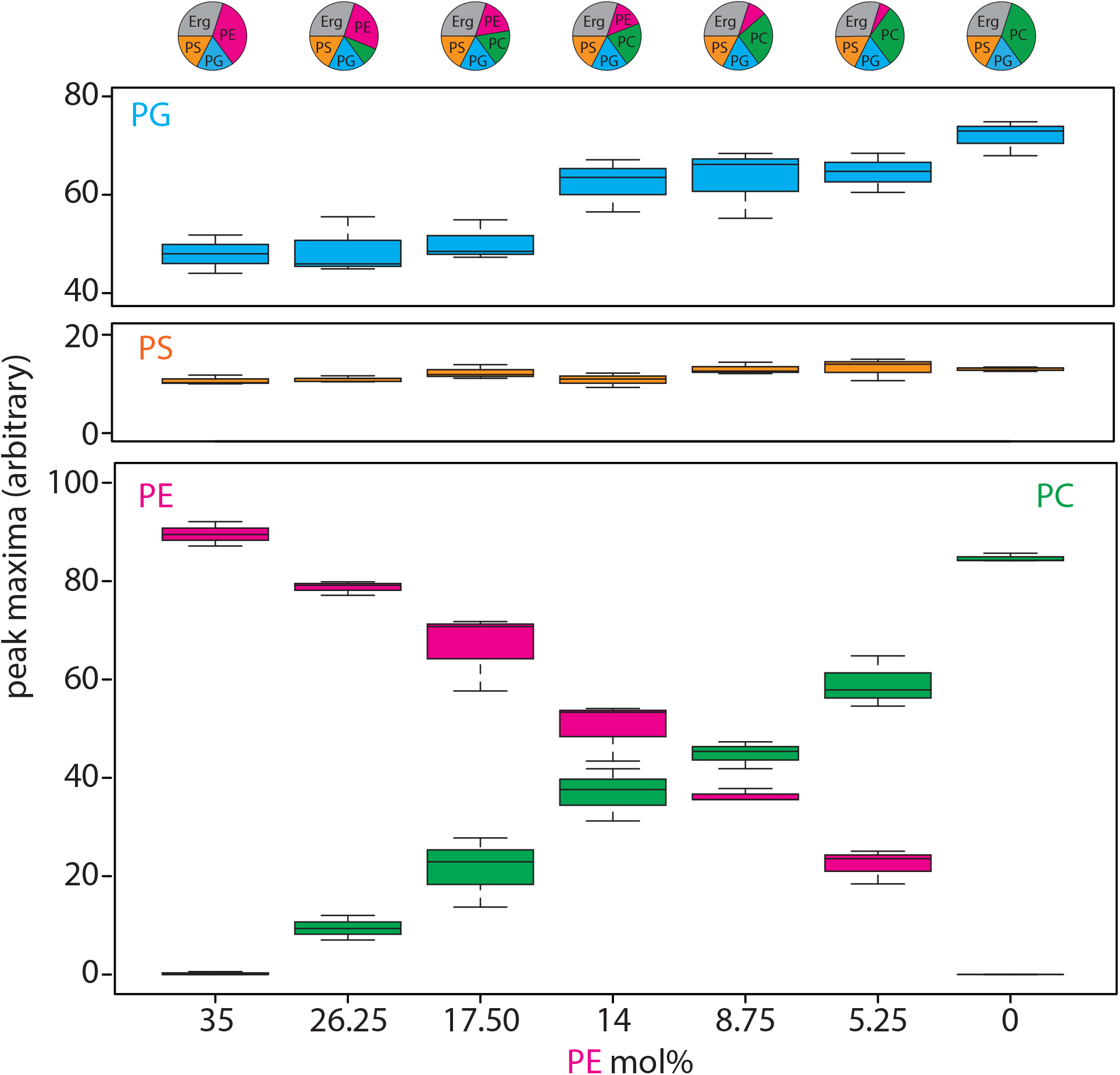
Lipid analysis by mass spectrometry of proteo-liposomes. Boxplots show peak areas of mass spectra corresponding to POPE, POPG, POPS and POPC, present in in respective Lyp1 vesicles. Absolute quantities of lipids (mol%) taken for each lipid composition are shown in the pie-graphs above. Two-letter abbreviation and color-coding: PC = phosphatidylcholine (green), PE = Phosphatidylethanolamine (magenta), PS = phosphatidylserine (orange) and PG = phosphatidylglycerol (blue). Line within boxplot represents the median. Top and bottom represent the first and third quartile, respectively. Error bars are the minimal and maximal value. Number of experiments = 3 experimental replicates.

**Figure S2.**
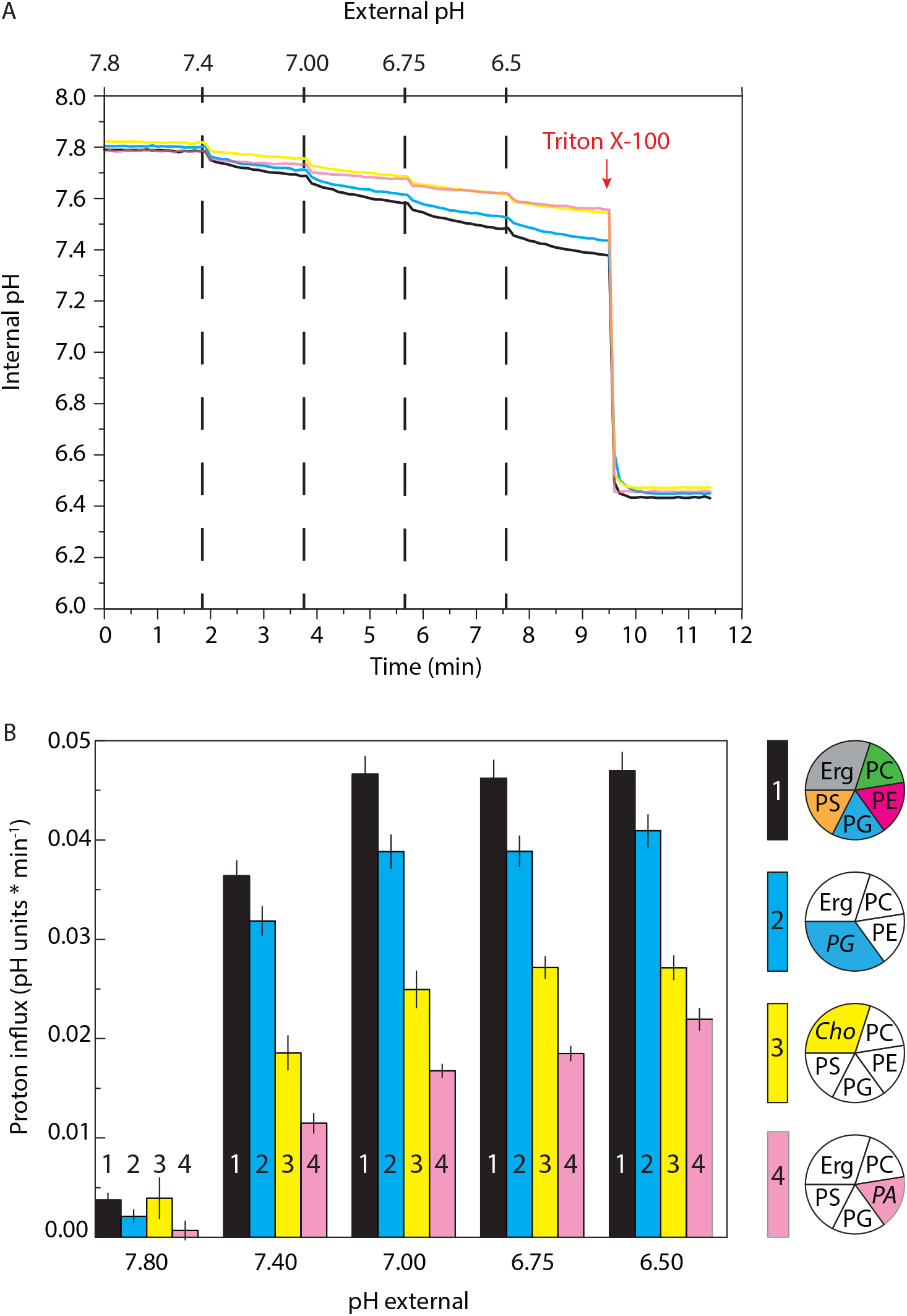
Proton permeability of proteoliposomes. (A) Internal pH is shown as a function of external pH and time for proteoliposomes composed of various lipids. The vesicles were prepared in 10 mM potassium-phosphate pH 7.8. Color code is: POPC/POPE/POPG/POPS/Ergosterol (17,5/17,5/17,5/17.5/30 mol%) = black; POPC/POPE/POPG/Ergosterol (17,5/17,5/35/30 mol%) = Blue; POPC/POPE/POPG/POPS/Cholesterol (17,5/17,5/17,5/17.5/30 mol%) = Yellow; POPC/POPG/POPS/POPA/Ergosterol (17,5/17,5/17,5/17.5/30 mol%) = Pink. The red-arrow indicates Triton X-100 addition, which solubilizes the proteoliposomes and releases pyranine. (B) Proton influx (pH units * min^−1^) is shown as a function of external pH for each lipid composition. The proton influx was calculated from the slope of a linear fit of the data, where dashed lines in Panel A indicate the start and end points used for the fit. Result is representative of three experiments, and error bars are the standard error of the fit.

